# Small-molecule cocktails induce the differentiation of human adipose-derived mesenchymal stem cells into hepatocyte-like cells

**DOI:** 10.1101/2021.07.19.452852

**Authors:** Kan Yin, Yang Xu, Di Wu, Weiyan Yang, Naijun Dong, Ning Li, Robert Chunhua Zhao

## Abstract

At present, liver transplantation and hepatocyte therapy are common methods for the treatment of end-stage liver diseases, but they are restricted due to the shortage of liver donors and the safety and effectiveness of hepatocyte sources. Human adipose-derived mesenchymal stem cells (HAD-MSCs) have been applied to efficiently and stably induce phenotypic and functional liver cells or tissues in vitro due to their advantages such as wide sources and easy access to materials. In this study, the HAD-MSCs liver differentiation induction system was established and optimized based on the “cocktail method” of chemical small molecule compounds. We used HAD-MSCs as seed cells and gradually obtained mature hepatoid cells with normal phenotype and function after induction with small molecule compounds and growth factor system in vitro. The hepatoid cells induced by the two groups showed high similarity in phenotype and functional characteristics of mature hepatocytes. The differentiation system of human adipose mesenchymal stem cells into hepatocytes induced by small-molecule compounds in vitro was successfully constructed. This study will lay a foundation for the optimization of liver differentiation strategies and provide a reliable source of functional liver cells for clinical studies of liver diseases.

## Introduction

Cellular identity in a broad sense can be understood as the spatiotemporal expression patterns of genes in cells. The understanding of the molecular basis of cell fate transformation which means cellular identity switch and its precise manipulation have always been the core issues of stem cell research[1]. The understanding of the mechanism of cell lineage development is the core clue guiding the differentiation of stem cells in vitro[2]. In the study of pluripotent stem cell differentiation, the success or failure of differentiation depends on the ability to effectively simulate in vivo development of specific cell lineages[3]. Cellular reprogramming using cocktails of transcription factors (TFs) affirms the epigenetic and developmental plasticity of mammalian cells[4]. Since the basic biological behavior of stem cells such as self-renewal and differentiation is a highly time-space dependent regulatory process, the flexibility and high controllability of chemical biological methods are particularly suitable for stem cell research[5]. Recently, the use of small molecules has emerged as a powerful tool to induce cell fate transition for their superior stability, safety, cell permeability, and cost-effectiveness[6]. Small molecules can play an important role in regulating cell fate, not only can regulate specific target signal and compared with gene regulation has obvious advantages: 1. the effect of small molecules is rapid and reversible, can change their concentration and combined use of different kinds of small molecules for regulation of protein function accurately and timing; 2.Small molecular compounds can be added or removed at any time to initiate or interrupt a specific action; 3.Synthetic chemistry because of its structural and functional diversity gives small molecules unlimited potential to accurately control the recognition and interaction between molecules[7].

According to the classical development biology theory, at the early stage of embryonic development, the zygote first develops into blastocyst, and then the blastocyst develops into outer, middle, and inner layers. The ectoderm eventually develops into tissues such as nerves and skin, the mesoderm into tissues such as the heart, blood, muscles and bones, and the endoderm into internal organs such as the lungs, liver, pancreas and intestine[8]. Human adipose tissue-derived mesenchymal stem cells (hAD-MSCs) were successfully induced to differentiate into adipocytes, osteoblasts, chondrocytes, skeletal muscle cells and other mesodermal cells, and they could also differentiate into other layer cells, proving that hAD-MSCs are pluripotent stem cells[9].

Hepatocyte generation is gathering interest with industrial and clinical parties due to its relevance in the areas of drug development, cell therapy, and disease modeling[10]. We demonstrate here the efficient generation of hAD-MSCs-derived hepatocyte-like cells (HLCs) through the use of pure small molecules cocktails strategy, removing the longstanding reliance on recombinant growth factors, which provides a cost-effective platform for in vitro studies of the molecular mechanisms of human liver development and holds significant potential for future clinical applications.

## Methods and Materials

### Human adipose tissue

This study was ethically approved by Ethics Committee of Affiliated Hospital of Qingdao University. All adipose tissue samples were collected from patients who underwent surgical liposuction at Affiliated Hospital of Qingdao University. All the participants provided written consent for this study.

### Culture of human mesenchymal stem cells

Human adipose tissue was obtained from patients undergoing liposuction. Fresh liposuction tissue was collected, digested, isolated and cultured according to the established method. Cells were cultured with M-ADSC culture medium containing Dulbecco’s modified Eagle medium (DMEM) DMEM/F-12, MCDB-201, 2% fetal bovine serum (FBS), 1 × insulin transferrin selenium, 10^−8^ M dexamethasone,10^−4^ M ascorbic acid 2-phosphate, 10 ng/ml EGF, 10 ng/ml PDGF-BB, and 1 ng/ml Activin A[32]. The normal human liver cell line L02 were purchased from the Chinese Academy of Medical Sciences and cultured in in RPMI-1640 Medium supplemented with 10% (v/v) fetal bovine serum according to the instructions and protocols. The medium was changed every 2-3 days, and the cells were passaged for subculturing or subsequent experiments when they grew to 80% confluence.

### Flow Cytometry, FCM

For flow cytometry analyses, cells were permeabilized with Cytofix/Cytoperm Fixation/Permeabilization kit for 15 minutes, direct-standard antibodies of CD34, CD105, CD29, CD31, CD44, CD73, CD90 and HLA-DR were incubated at 4°C in dark for 30min. After incubation, cells were washed three times and analyzed by the BD Accuri C6. Data were analyzed with CFlow sample analysis software.

### Lipogenic and Osteogenic induction and identification of hAD-MSCs

After the cell density increased to 80-90%, the lipid induction medium was replaced. Oil red O staining was performed at 3 weeks after induction. The formation of lipid droplets was observed under the microscope and photographs were taken. Alkaline phosphatase staining and alizarin red staining were performed on the 9th day of induction. The blue-purple precipitation and calcium nodular precipitation were observed under the microscope.

### Hepatocyte differentiation in vitro

The hAD-MSCs in the small molecule compound (SM) group were first induced to Definitive Endoderm (DE): Serum free F12-DMEM medium containing 0.5% DMSO was added to induced for 24h. 3M CHIR99021 serum-free RPMI-1640 medium was added and cultured for 48h. Then serum-free RPMI-1640 medium was added and induced for 48h. Hepatoblasts were induced from DE cells: Add serum-free F12-DMEM medium containing 0.5M SB431542, 250nM SB(Sodium butyrate) and 0.5%DMSO for 8 days. Obtain Hepatocyte-like cells (HLCs): Serum-free F12-DMEM medium containing 15 M FH1, 15 M FPH1, 0.5 M SB431542, 100nM Dexamethasone, and 10 M Hydrocortisone was added and the medium were changed daily for 6 days. In the growth factor (GF) group, low-serum (0.5% FBS) H-DMEM medium containing 5ng/ml Activin A and 50ng/ml Wnt3a was added and induced for 24h. H-DMEM medium with low serum (0.5% FBS) containing only 5ng/ml Activin A was added and induced for 4 days. Low-serum (1% FBS) H-DMEM medium containing 10ng/ mL FGF-BASIC and 10ng/ml BMP-4 was added and induced for 4 days. Low-serum (0.5% FBS) H-DMEM medium containing 10ng/mL FGF-4 and 10ng/ml HGF was added for 4 days. Low-serum (0.5% FBS) H-DMEM medium containing 10ng/mL FGF-4, 10ng/ml HGF, 10ng/ml OSM and 100nM DEX was added and induced for 6 days.

### Real-time-PCR

Total RNA was extracted using purification Kit. cDNA synthesis was performed with 1μg of RNA using PrimeScript reverse transcriptase and reverse transcribed using a PCR Instrumentation C1000 Touch(tm) Thermal Cycler (Bio-Rad Laboratories, Hercules, CA, USA) according to the manufacturers’ protocol. RT-PCR analysis was performed on an ABI Prism 7500 Sequence Detection System using the SYBR Green PCR Master Mix. The gene expression of DE markers [sex determining region Y (SRY)-box 17 (SOX17), FOXA2], hepatic progenitor markers [alpha-fetoprotein (AFP), HNF4α, cytokeratin 18 (CK18), cytokeratin 19 (CK19)], and hepatocyte markers [albumin (ALB), alpha-1 antitrypsin (A1AT), CYP1A2, CYP3A4] were measured. GAPDH expression was used as an internal control.

### Immunofluorescence microscopy

Cells at each differentiation stage were fixed with iced methanol or 4%paraformaldehyde for 15 min at room temperature and blocked with phosphate-buffered saline (PBS) containing 0.1% Triton X-100 and 3% bovine serum albumin (BSA) at room temperature for 1 h. Cells were then incubated with the appropriate primary antibodies at 4 °C overnight. On the second day, after three washes for at least 5 min with PBS, Alexa Fluor conjugated secondary antibody diluted 1:1000 was added and incubated at room temperature for 1 h. Hoechst 33342 diluted in 1:5000 was used to stain the cell nuclei. Between each step, cells or sections were washed with fresh PBS. Image acquisition and processing were carried out using a fluorescence microscope (Zeiss LSM 800, Germany). and counted by Image-Pro Plus software.

### Western blot analysis

Cells were lysed in ice-cold RIPA cell buffer supplemented with a protease inhibitors cocktail. After centrifuging at 12,000 rpm for 10 min at 4 °C, the supernatant was collected as the total cell lysate. The membrane was blocked with 5% nonfat milk for 1 h at room temperature, incubated overnight at 4 °C with the relevant primary antibodies, and then incubated with horseradish peroxidase-conjugated secondary antibodies for 1 h at room temperature. The information of indicated primary antibodies (anti-Foxa2 1:1000, anti-Sox17 1:5000, anti-HNF4α 1:1000, anti-AFP 1:4000, anti-ALB 1:20000, anti GAPDH 1:20000). The secondary antibodies used were horseradish peroxidase (HRP)-conjugated goat anti-mouse IgG and goat anti-rabbit IgG. Protein bands were visualized with ECL substrates.

### Albumin secretion ELISA assay

At the endpoint of the differentiation process, the supernatant of cultured cells was collected. Albumin secretion in the supernatant was measured with a human albumin enzyme-linked immunosorbent assay (ELISA) quantitation kit according to the manufacturer’s instructions. Cells were trypsinized and counted with Cell Counter. The albumin secretion was normalized to total cell numbers.

### Periodic acid-Schiff staining for glycogen

Periodic acid-Schiff (PAS) is a staining method that is primarily used to identify glycogen storage in cells. Cells were fixed in 4% paraformaldehyde and stained using a PAS staining system at room temperature. Briefly, fixed cells were oxidized with 1% periodic acid solution, then incubated in Schiff’s reagent. After being rinsed with PBS, cells were stained with Mayer’s hematoxylin. Between each step, the cells were washed with fresh PBS.

### Cytochrome P450 activity

CYP1A2 activity was measured using a CYP1A2-MROD Assays kit. The assay utilizes a nonfluorescent CYP1A2 substrate that is converted into a highly fluorescent metabolite (resorufin) detected in the visible range (Ex/Em = 530/590 nm). For CYP1A2 induction, omeprazole (100 μM) was added to the differentiated human ES and iPS cells during the last 3 days and human primary hepatocytes for 72 h. The medium was refreshed every day. The cells were lysed with RIPA and then homogenized with an ultrasonic crusher. The assay was performed according to the manufacturer’s instructions. The fluorescence was measured. Cytochrome activity was normalized to the total protein (mg) and presented as pmol/mg protein/min.

### Statistical analysis

All experiment was performed at least three independent times. Data are presented as the mean ± standard deviation (SD). GraphPad Prism 5 (USA) was used for all statistical analyses. Two-tailed Student’s t test was used for comparisons of two independent groups, and the level of statistical significance was set at P < 0.05 (*p < 0.05).

## Results

### Characterization of hAD-MSCs

Under an inverted microscope, hAD-MSCs grew mainly in the form of long flat fusiform adherent walls. After the cells grew to 80%∼90% fusion, the hAD-MSCs were cultured for subculture. The growth rate of the cells after passage was significantly faster than that of the primary cells, and the cells were arranged in a vortex pattern, and the cells could be passed on after 3-5 days of growth (Fig1A). Flow cytometry was used to identify the surface markers of the third-generation hAD-MSCs. The results showed that hAD-MSCs were highly expressed in CD29(99.6%), CD44(99.5%), CD73(99.9%), CD90(99.8%) and CD105(99.1%), while CD31(1.4%), CD34(0.4%) and HLA-DR(0.5%), indicating that the extracted cells contain the characteristics of hAD-MSCs (Fig1B). hAD-MSCs can differentiate into adipocytes and osteoblasts under the stimulation of specific medium. Under the continuous induction of adipogenic induction medium, the morphology of hAD-MSCs gradually changed from spindle shape to round shape, with lipid droplets appearing in the cytoplasm. After 3 weeks of induction, oil-red O staining was performed, indicating large red lipid droplets in the cytoplasm (Fig1B). Under the continuous induction of osteogenic induction medium, the cells appeared stacking arrangement and growth, and mineral salt deposition was visible to the naked eye. On the 9th day after induction, alkaline phosphatase and alizarin red staining were performed, and blue precipitates and calcium nodules were observed microscopically (Fig1B). Alkaline phosphatase (ALP) is the marker of mature osteoblasts, and calcium nodules are also markers of osteoblasts.

**Fig1.**
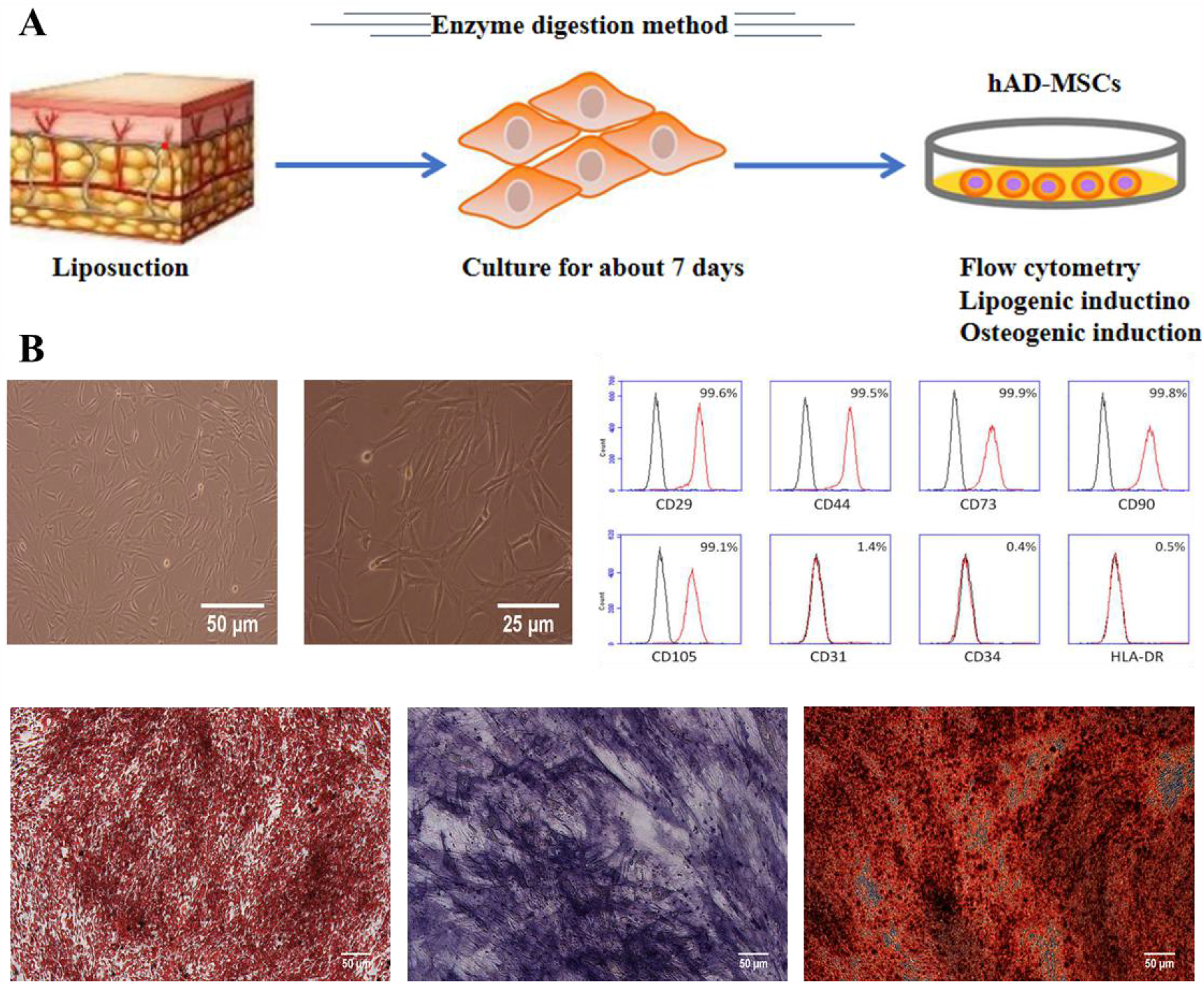
Characterization of hAD-MSCs. A. Isolation, extraction and culture of primary hAD-MSCs. B. hAD-MSCwere mainly long-flat fusiform; were highly expressed in CD29, CD44, CD73, CD90 and CD105, while lowly in CD31, CD34 and HLA-DR; showed adipogenic and osteogenic differentiation ability with oil-red O, alkaline phosphatase and alizarin red staining. Scale bars = 50 μm.

### Morphological changes during hepatoid cell differentiation induced in vitro

We have therefore developed a differentiation procedure that is devoid of growth factors and driven by small molecules. This stepwise approach is able to generate HLCs at high efficiency. The procedure is notionally trisected into three phases inducing DE differentiation (phase I), hepatic specification (phase II), and hepatocyte maturation (phase III). Similar experiments were also performed with growth-factor-based approaches and consistent results were obtained (Fig2A). Under the periodic stimulation of small molecular compounds and growth factors, on the 5th day of induced differentiation, the cells were observed to increase in size under the microscope, and the cells changed from long-spindle to petal cluster. On the 9th day of induced differentiation, the intercellular space increased and the number of cells decreased. On the 13th day, the cells were irregular or square, and the number of cells increased. At the end of differentiation on the 19th day of the final stage, the microscopic appearance of the cells was similar to that of hepatocytes (Fig2B).

**Fig2.**
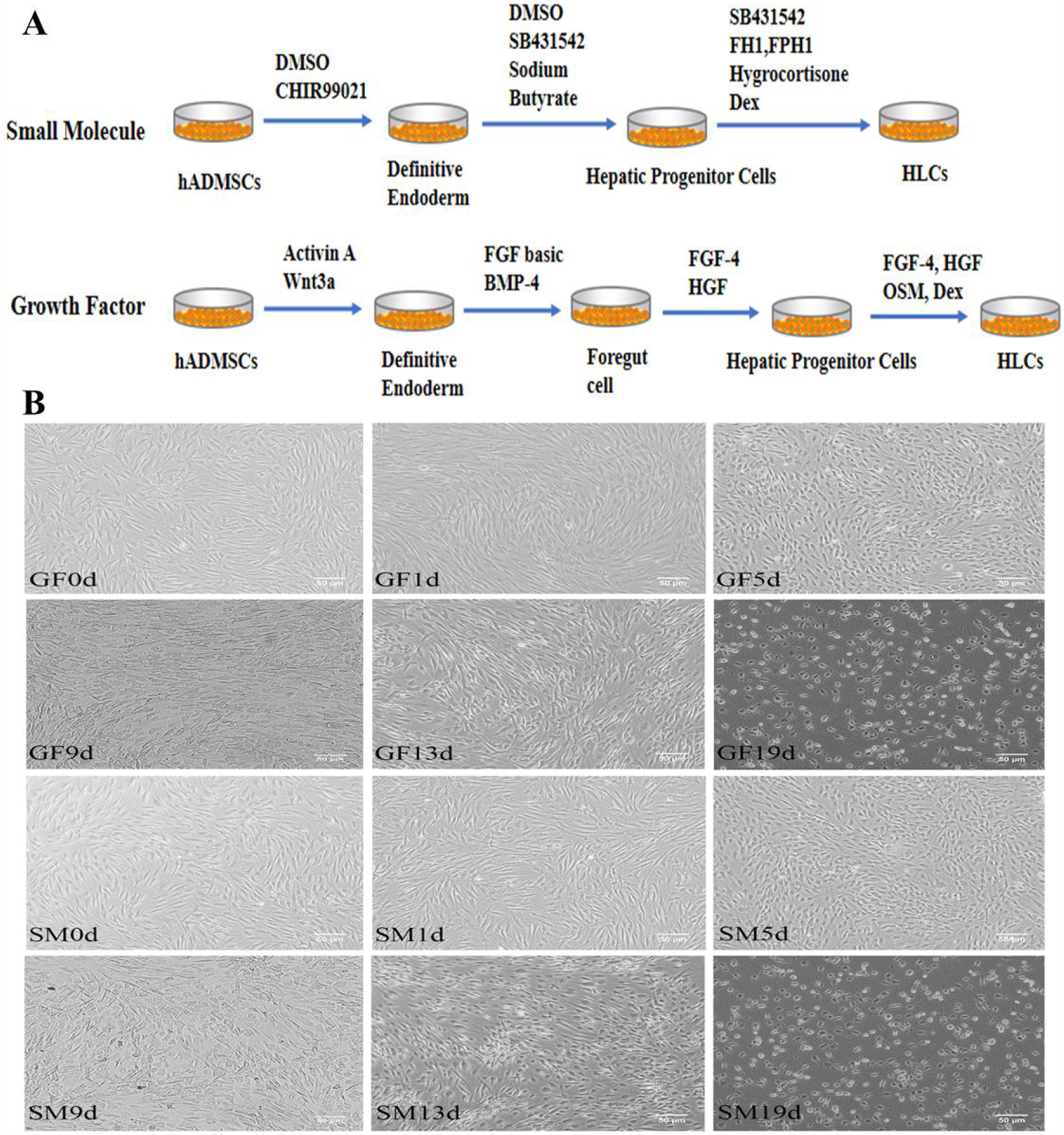
Schematic diagram of induced hAD-MSCs differentiation into hepatocytes. A. The detailed hepatocyte differentiation protocol using pure small molecules or growth factors. B. Sequential morphologic changes during the differentiation process. Scale bars = 50 μm.

### Phase I: Production of definitive endoderm(DE)

We set out to investigate whether CHIR99021 (CHIR), an inhibitor of GSK3β which can indirectly activate Wnt/β-catenin signaling, could promote definitive endoderm differentiation from hAD-MSCs. 3μM CHIR99021 was added to serum-free RPMI-1640 medium in small molecular group(SM), at the same time, in growth factor group(GF) we use 5ng/ml Activin A and 50ng/ml Wnt3a added in low serum (containing 0.5% FBS) H-DMEM medium. Following treated procedure, at day5, we observed elevated expression of DE markers such as Sox17, Foxa2, CXCR4 and GATA4. With qRT-PCR and immunofluorecence. The undifferentiated hAD-MSCs and L-02 cells were used as controls (Fig3).

**Fig3.**
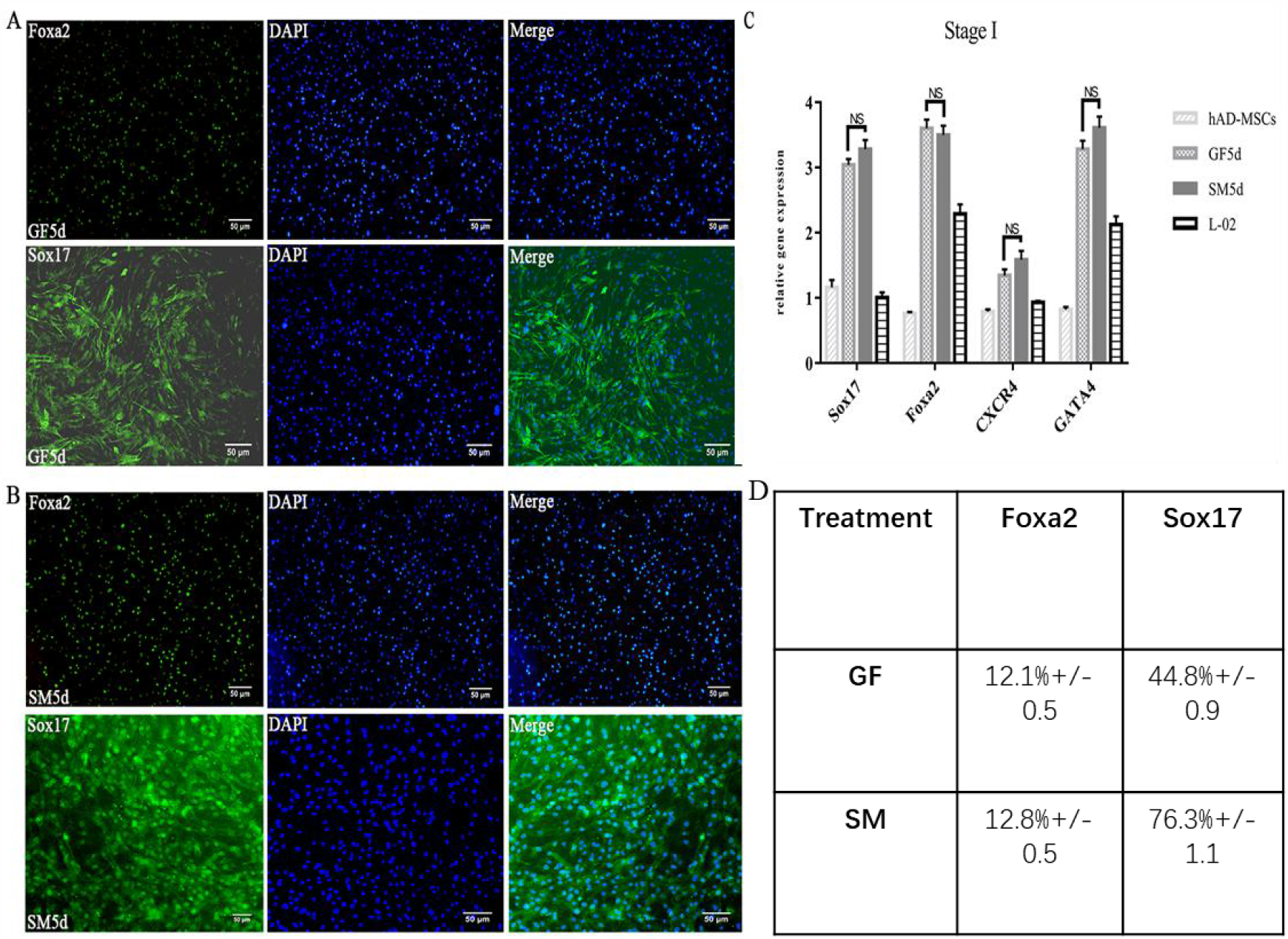
Small molecules efficiently induce definitive endoderm differentiation. A. Expression of Foxa2 and Sox17 at protocol endpoints using growth factors. B. Expression of Foxa2 and Sox17 at protocol endpoints using small molecules. C.q-PCR analysis of markers (Sox17,Foxa2,CXCR4 and GATA4) from small-molecule cocktails or growth factors. D. Efficiency of phase I differentiation, determined by counting Fox2-positive cells, and Sox17 positive cells. Efficiencies are presented as the percentage of positive cells plus or minus the SD of all fields counted. Scale bars = 50 μm.

### Phase II: Hepatic Specification

To specify a hepatic fate, we explored potential routes to effificiently produce an AFP/HNF4A-positive hepatic progenitor population. We used a well-established method consisting of serum-free F12-DMEM medium based on 0.5 M SB431542, 250 nM SB(Sodium butyrate) and 0.5%DMSO in SM group, low-serum (1% FBS) H-DMEM medium containing 10ng/ mL FGF-BASIC and 10ng/ml BMP-4 to induce forgut cells, then low-serum (0.5% FBS)H-DMEM medium containing 10ng/ mL FGF-4 and 10ng/ml HGF to induce hepatoblasts in GF group. Gene expression of hepatic progenitor markers were analyzed after the end of hepatic specifification by qRT-PCR and immunofluorecence. There was no significant difference in the expression of ALB, AFP, HNF4, CK18and CK19 marker genes between SM group and GF group, while CK18 expression in SM group was significantly higher than that in GF group (Fig4C).

**Fig4.**
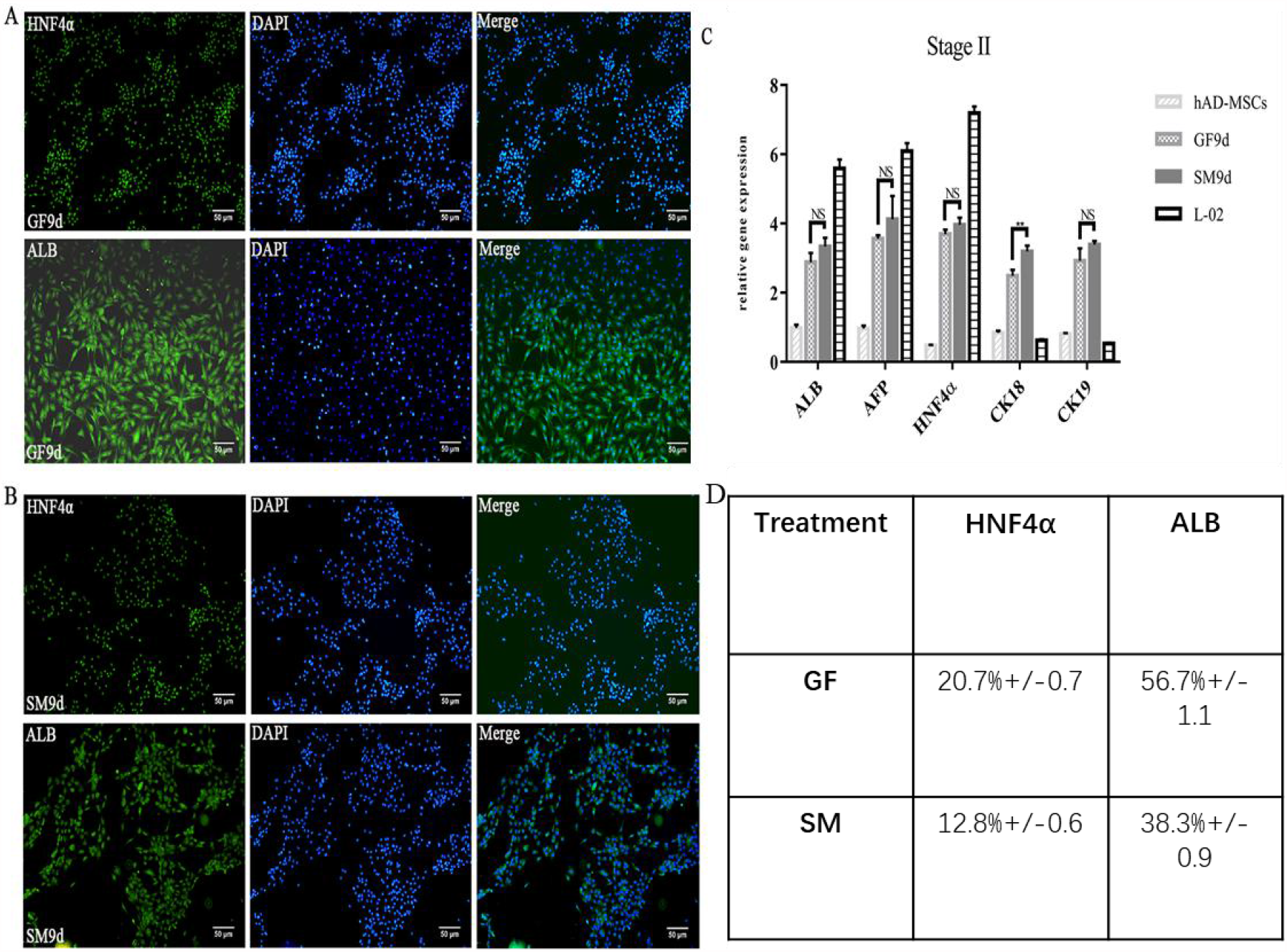
Small molecules efficiently induce hepatic progenitors from definitive endoderm. A. Expression of HNF4α and ALB at protocol endpoints using growth factors. B. Expression of HNF4α and ALB at protocol endpoints using small molecules. C. q-PCR analysis of markers (ALB,AFP, HNF4α,CK18and CK19) from small-molecule cocktails or growth factors. D. Efficiency of phase I differentiation, determined by counting HNF4α-positive cells, and ALB-positive cells. Efficiencies are presented as the percentage of positive cells plus or minus the SD of all fields counted. Scale bars = 50 μm.

### Phase III: hepatic maturation, from hepatoblasts to hepatocyte like cells(HLCs)

In well established in hepatocyte maturation procedures, serum-free F12-DMEM medium containing 15 M FH1, 15 M FPH1, 0.5 M SB431542, 100nM Dexamethasone and 10 M Hydrocortisone in SM group, low serum (0.5% FBS) H-DMEM medium containing 10ng/ mL FGF-4, 10ng/ml HGF, 10ng/ml OSM and 100nM Dexamethasone in GF group. We observed comparable effificiencies of differentiation between the SM and GF approaches by assessing ALB, AFP, A1AT, CYP1A2 and CYP3A4, the A1AT expression in SM group was significantly higher than that in GF group (Fig5). In conclusion there were higher levels of expression of all hepatic markers in hepatocyte like cells (HLCs) in 2 types of treatment.

**Fig5.**
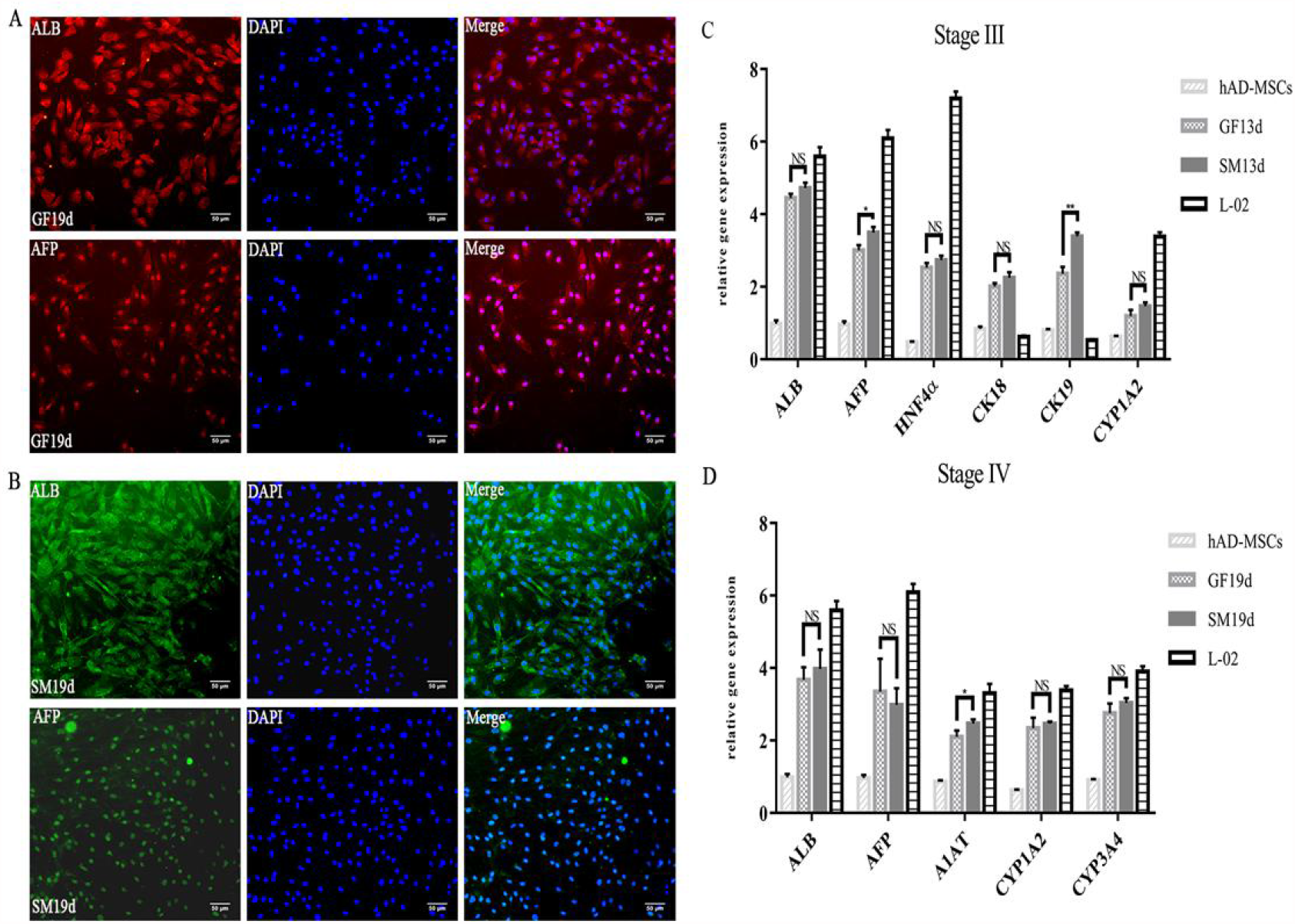
Characterization of phase III and IV differentiation to hepatocyte-like Cells. A. Expression of AFP and ALB at protocol endpoints using growth factors. B. Expression of AFP and ALB at protocol endpoints using small molecules. C. q-PCR analysis of markers (ALB,AFP, HNF4α,CK18, CK19 and CYP1A2) from small-molecule cocktails or growth factors at 13 day. D. q-PCR analysis of markers (ALB,AFP, A1AT,CYP1A2 and CYP3A4) from small-molecule cocktails or growth factors at 19 day. Scale bars = 50 μm.

### Small-Molecule-Derived HLCs Demonstrate Hepatic Function

Liver was mainly used for synthetic secretion and metabolism function in organs, in order to test our add in stages of small molecules and growth factors induced cells have gained by the functional characteristics of mature hepatocytes, albumin secretion to experiment, the results showed that GF and SM groups all have the ability to secrete albumin and no significant difference between two groups. Western blot was used to determine the total expression of marker gene ALB at the protein level. Western blot results showed that the expression of albumin in GF and SM groups was positively correlated with the maturity of liver cells. The content of albumin in the supernatant of SM group was about 12 g/ mL, slightly higher than that of GF group (Fig6A). Glycogen storage is another important feature of functional hepatocytes. The ability to store glycogen was measured by adding medium with a sugar concentration of 450mg/dl. Experimental data showed that GF and SM groups had the ability to store glycogen (Fig6C), similar to functional liver cells. In order to test whether hepatocytes from GF group and SM group have the ability to convert metabolic drugs and isobiomass, we evaluated their cytochrome enzyme activity. CYP1A2 is one of the major enzymes in drug metabolism in human. After treatment with 10 M omeprazole, liver cells induced by GF and SM groups showed the ability to metabolize drugs (Fig6B). These liver specific function tests were consistent with the expression of mature hepatocyte markers, suggesting that the induction differentiation system based on small molecules and growth factors could induce hADMSCs to differentiate into functional hepatocytes in vitro. There was no significant difference in liver function expression between hepatocytes obtained from GF group and SM group.

**Fig6.**
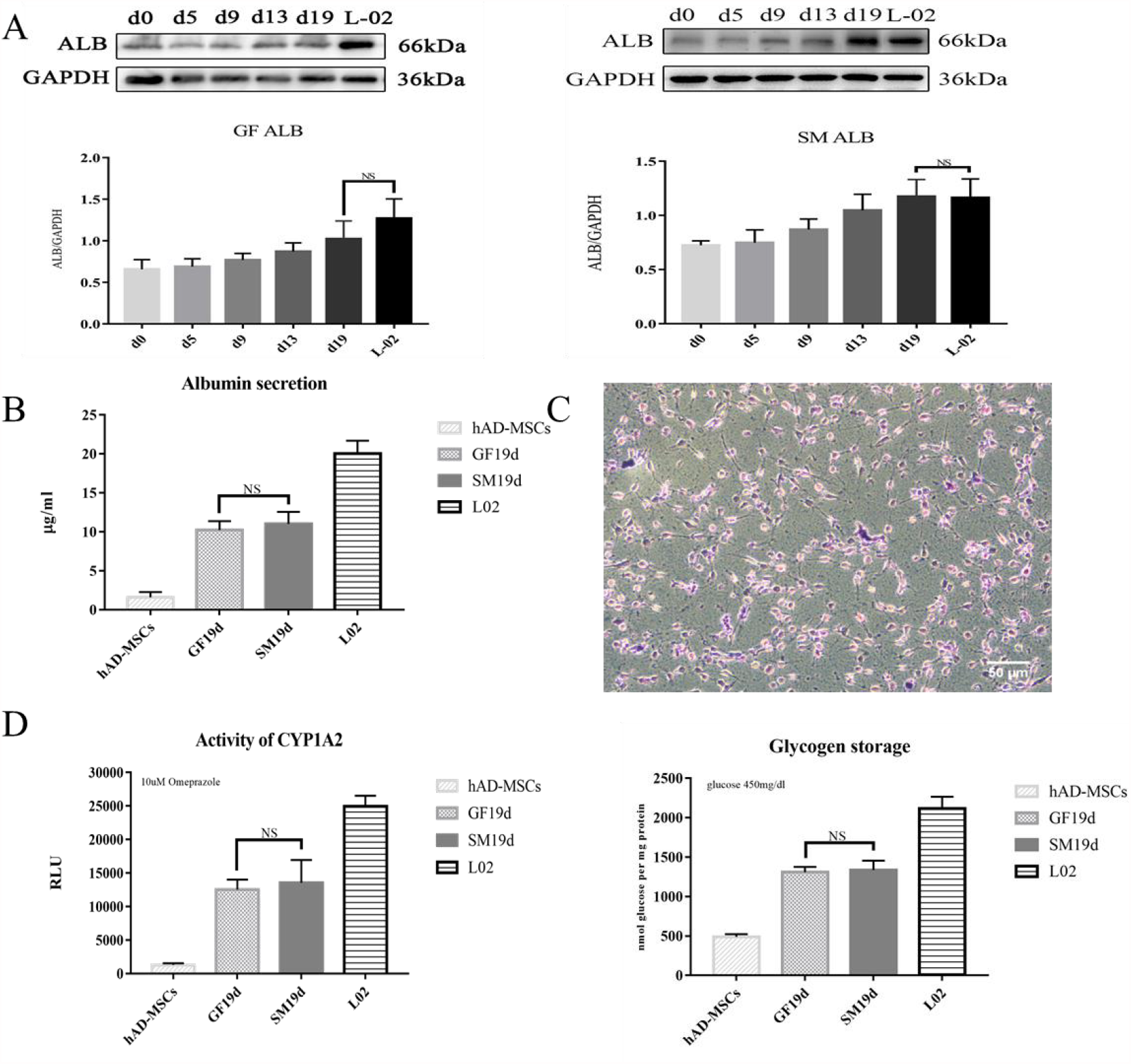
Functional analysis of SM and GF induced hepatocyte-like cells. A. Western blotting for albumin expression of in SM-iHep and GF-iHep. B. Albumin secretion of the differentiated cells treated with small-molecule cocktails or growth factors. C. PAS staining showing glycogen storage in SM- and GF-induced differentiated cells. L02 hepatocytes were used as the control (*p value < 0.05, **p value < 0.01). D. Cytochrome P450 1A2 activity in SM-iHep and GF-iHep after induction with omeprazole.

## Discussion

In this study, we first used hAD-MSCs to obtain the limited endoderm cells by adding small molecular compounds and growth factors in vitro, and then identified the limited endoderm cells at the gene and protein levels, mature liver cells expressing liver-specific markers were obtained. The establishment and optimization of the liver differentiation system of adult stem cells with small molecule compounds lay a foundation for the study of the molecular mechanism of human liver development in vitro and provide a new strategy for clinical and pharmacological applications in the future. The liver is the largest substantive visceral organ in the human body, which plays the role of storing liver glycogen, synthesizing secretory protein, regulating lipid metabolism, etc.[11]. The liver is also the production site of bile, has the function of detoxification, and participates in the immune response of the body[12]. It is an indispensable and important visceral organ in the body. The molecular mechanism of liver regeneration is complex and orderly, and many gene expressions are significantly up-regulated, involving the synergistic effect of hundreds of genes or proteins, regulated by factors including growth factors, cytokines, transcription factors (TFs), hormones, oxidative stress products, metabolic networks, and microRNA[13]. How to explain the molecular mechanism of liver regeneration deeply from the biological phenomenon of liver regeneration is a great challenge. Most of the researches on early liver regeneration are confined to the tissue and cell level, which are studied from the aspects of tissue morphology and cell structure, but the research on the regulation mechanism at the molecular level is still insufficient[14]. With the development of high-throughput detection technology such as gene chip, the mechanism of liver regeneration is deeply studied through omics tools such as genome, proteome and metabolome, in an attempt to reveal the precise regulation mechanism of liver regeneration from the systematic level[15].

Liver transplantation and hepatocyte therapy are effective means to treat irreversible liver diseases. However, all these treatments are limited by organ or cell resources, so developing enough functional liver cells for liver regeneration is imperative[16]. The Nodal signaling pathway is the core of the limiting endoderm stage, and activation of the Nodal/TGF-signaling pathway is triggered by TGF-family member Activin A, leading to phosphorylation of SMAD2 and SMAD3, which are then bound to SMAD4, transported to the nucleus, and regulated transcription genes mediating cell proliferation or differentiation[17]. Then FGF and BMP signals continue to regulate the development of the liver and gradually enter the stage of hepatocyte specialization[18].Transcription factors FoxA and GATA families, as well as Hhex, are involved in early liver development and determine liver development[19].With the expression of liver enrichment transcription factors such as HNF4 and C/EBP, embryonic liver began to form and gradually expressed specific marker genes of liver cells, albumin (ALB), alpha Fetoprotein (AFP), keratin families CK18 and CK19, etc.[20,21].

In this study, we established a highly efficient, reproducible and efficient induction system by referring to published methods of MSCs differentiation into liver cells. Most induction schemes use a phased induction method to simulate the development of liver in vivo, in which hAD-MSCs are first induced to form limited endoderm cells, then to differentiate into hepatic progenitor cells, and finally to induce mature liver cells. At the induction and restriction endoderm stage, two different dedifferentiation methods are used: activins A and Wnt3a growth factor and small molecules CHIR99021, A GSK3 inhibitor. These two dedifferentiation methods have been reported in previous studies[22,23]. However, when applied to hAD-MSCs, we noted that both of the previously published approaches required some degree of fine-tuning to achieve efficient induction. Previous studies have found that DMSO and sodium butyrate (SB) can guide the differentiation of DE into liver lineage[24,25].Restricted endodermal differentiation during liver development usually produces a range of developmental outcomes, including hepatocytes, pancreatic cells, and intestinal cells, etc., which inhibit TGF-pathway to differentiate restricted endodermal cells into hepatic progenitor cells[6]. SB431542 is an effective selective ALK5 inhibitor, combined with sodium butyrate and DMSO, which can promote the DE stage liver specialization of SM group cells. It has been reported that small molecule compounds FH1 and FPH1 can induce proliferation and enhance the function of primary human hepatocytes cultured in vitro[26]. These two small molecule compounds, used in combination with dexamethasone and hydrocortisone, are applied in the final stage of hepatocyte production by hepatic progenitor cells. Mature hepatocyte-like cells were obtained in SM group, which not only expressed AFP, ALB, HNF4 and other marker genes, but also showed functions of albumin secretion, glycogen storage and drug metabolism. In GF group, mature liver cells were obtained by four-step induction[10], and compared with relevant differentiation stages in SM group at the end of each stage of differentiation. The data showed that the mature liver cells induced by SM group and GF group were consistent in both the expression of liver-specific genes and the function of mature liver cells, with no significant difference. In summary, we demonstrate the high efficiency of hepatocyte-like production by combining multiple small molecule compounds to reduce long-term dependence on recombinant growth factors. This may provide the basis for mass production of hepatocytes in vitro and has important clinical application potential.

With the optimization and improvement of differentiation strategies, most of the hepatoid cells generated by published research schemes have shown similar functional results with primary hepatocytes in vitro, and even can proliferate and perform corresponding functions in the liver after the induced liver cells are transplanted into animal models[27,28].Until now, however, the maturity of most differentiated cells has yet to be compared with their counterparts in the body. Small molecular compounds have made rapid progress in regulating the fate of cells. Many cell types, such as neurons, cardiomyocytes, and retinal pigment epithelial cells, have been generated by the sequential addition of specific small molecular compounds to PSCs by simulating organ development in vivo[29].Meanwhile, small molecular compounds have been successfully used for reprogramming and transdifferentiation, etc.[5,30].In recent years, scientists have found that fibroblasts are transdifferentiated into liver cells after the induction of small molecule compounds and growth factors[31].In the future, more small molecular compounds are waiting to be discovered to replace growth factors. While future applications of these small molecules to generate cells should be considered, more research should be done to improve their function and precisely control their destiny differentiation.

## Conclusions

After the above research and analysis, we come to the following conclusion. In this study, through periodic addition of different small molecule compounds and growth factors in vitro, mature hepatocytes with good phenotype and expression of mature hepatocyte function were obtained, and a small molecule compound induction system for hAD-MSCs differentiation into hepatocytes was successfully established. It provides an economical and effective platform for studying the molecular mechanism of human liver development in vitro.

